# The impact of YabG mutations on *C. difficile* spore germination and processing of spore substrates

**DOI:** 10.1101/2024.06.10.598338

**Authors:** Morgan S. Osborne, Joshua N. Brehm, Carmen Olivença, Alicia M. Cochran, Mónica Serrano, Adriano O. Henriques, Joseph A. Sorg

## Abstract

YabG is a sporulation-specific protease that is conserved among sporulating bacteria. *C. difficile* YabG processes cortex destined proteins preproSleC into proSleC and CspBA to CspB and CspA. YabG also affects synthesis of spore coat/exosporium proteins CotA and CdeM. In prior work that identified CspA as the co-germinant receptor, mutations in *yabG* were found which altered the co-germinants required to initiate spore germination. To understand how these mutations in the *yabG* locus contribute to *C. difficile* spore germination, we introduced these mutations into an isogenic background. Spores derived from *C. difficile yabG*_C207A_ (catalytically inactive), *C. difficile yabG*_A46D_, *C. difficile yabG*_G37E,_ and *C. difficile yabG*_P153L_ strains germinated in response to TA alone. Recombinantly expressed and purified preproSleC incubated with *E. coli* lysate expressing wild type YabG resulted in the removal of the pre sequence from preproSleC. Interestingly, only YabG_A46D_ showed any activity towards purified preproSleC. Mutation of the YabG processing site in preproSleC (R119A) led to YabG shifting its processing to R115 or R112. Finally, changes in *yabG* expression under the mutant promoters were analyzed using a SNAP-tag and revealed expression differences at early and late stages of sporulation. Overall, our results support and expand upon the hypothesis that YabG is important for germination and spore assembly and, upon mutation of the processing site, can shift where it cleaves substrates.

## Introduction

Susceptibility to *Clostridioides difficile* infection (CDI) is commonly associated with the use of broad-spectrum antibiotics that disrupt the normally protective colonic microbiota (Theriot *et al*., 2014, Buffie *et al*., 2015, Di Bella *et al*., 2024). Subsequent ingestion of *C. difficile* spores results in the germination of the spore form to the toxin-producing vegetative form (Paredes-Sabja *et al*., 2014). *C. difficile* vegetative cells secrete two toxins that damage the colonic epithelium – resulting in the symptoms associated with CDI (*e.g.,* diarrhea or pseudomembranous colitis) (Smits *et al*., 2016). Because spores are metabolically dormant and resist antibiotic action / harsh environmental conditions, they are the dispersive and transmissive form of the organism (Deakin *et al*., 2012). Unfortunately, the primary treatment for CDI is additional broad-spectrum antibiotics (*e.g.,* vancomycin or fidaxomicin) (Zar *et al*., 2007). Though these antibiotics treat the toxin-producing vegetative cells, but not the dormant spores, they continue to disrupt the colonic microbiota and can leave the patient susceptible to reinfections (Smits *et al*., 2016). Approximately 15-20% of patients experience recurring disease, and the likelihood of subsequent episodes increases with each recurrence (Di Bella *et al*., 2024).

*C. difficile* spores germinate in response to certain host-derived bile acids (*e.g.,* taurocholic acid) and amino acids (*e.g.,* glycine) (Shrestha & Sorg, 2018, Sorg & Sonenshein, 2009, Sorg & Sonenshein, 2008, Howerton *et al*., 2011, Ramirez *et al*., 2010, Bhattacharjee *et al*., 2016, Shrestha *et al*., 2017, Wilson, 1983, Wilson *et al*., 1982). Prior work provided genetic evidence that CspC is the bile acid germinant receptor and CspA is the co-germinant (amino acid) receptor (Francis *et al*., 2013, Shrestha *et al*., 2019, Kevorkian & Shen, 2017, Rohlfing *et al*., 2019). Despite the absence of a catalytic triad in these subtilisin-like proteases, these two pseudoproteases regulate *C. difficile* spore germination (Kevorkian & Shen, 2017, Kevorkian *et al*., 2016, Shrestha *et al*., 2019, Francis *et al*., 2013). Interestingly, *cspA* is encoded as a translational fusion to *cspB* (*cspBA*) and *cspC* is encoded downstream in the same operon (Paredes-Sabja *et al*., 2011, Bhattacharjee *et al*., 2016, Adams *et al*., 2013). CspBA is post-translationally processed to CspB and CspA by the sporulation-specific protease, YabG (Shrestha *et al*., 2019, Kevorkian *et al*., 2016, Kevorkian & Shen, 2017). In addition to processing CspBA, YabG also cleaves the pre-sequence from preproSleC (a cortex degrading enzyme) to generate proSleC, the form present in mature spores (Shrestha *et al*., 2019, Kevorkian *et al*., 2016). We hypothesize that CspB is associated with CspC and CspA in dormant spores, with CspC and CspA inhibiting the proteolytic activity of CspB. Upon binding of the bile acid and co-germinant, CspC and CspA dissociate from CspB. CspB then cleaves the inhibitory pro-peptide from proSleC, resulting in activation of SleC, cortex degradation, and the subsequent steps in germination (Shrestha *et al*., 2019, Zhu *et al*., 2018, Adams *et al*., 2013, Francis *et al*., 2015). The absence of *yabG* has also been shown to influence the extractability/expression of coat and exosporium proteins, SpoIVA, CotA, CotE, and CdeM in a *C. difficile yabG* mutant, suggesting other targets of YabG (Marini *et al*., 2023, Zhu *et al*., 2018).

YabG is a conserved sporulation specific protease. In *Bacillus subtilis*, YabG is important for proper assembly of the spore coat and has been shown to process at least six coat proteins (SpoIVA, CotF, CotT, YeeK, YxeA, and SafA) (Takamatsu *et al*., 2000a, Takamatsu *et al*., 2000b). Of those proteins, only SpoIVA has an orthologue in *C. difficile* and may be a YabG substrate (Kevorkian *et al*., 2016). YabG is produced during sporulation in the mother cell, under the alternative RNA polymerase sigma factors σ^E^ and σ^K^ (Fimlaid *et al*., 2013, Pereira *et al*., 2013, Saujet *et al*., 2013, Takamatsu *et al*., 2000b). *C. difficile* YabG is required for the expression of two σ^K^-dependent genes *cotA* and *cdeM*, which encode a coat and an exosporium protein, respectively (Marini *et al*., 2023).

Using an ethyl methane sulfonate (EMS) screen to identify strains which germinated without a co-germinant, we identified mutations in the *yabG* coding and promoter regions (Figure 1 and Table S3) (Shrestha *et al*., 2019). Moreover, these *yabG* mutant strains incorporated preproSleC and CspBA (the unprocessed forms) into spores. One of the *yabG* EMS mutant alleles, *yabG*_A46D_, (Table S3) appeared to have lower activity than wildtype, but the data was inconclusive (Shrestha *et al*., 2019).

**Figure 1.**
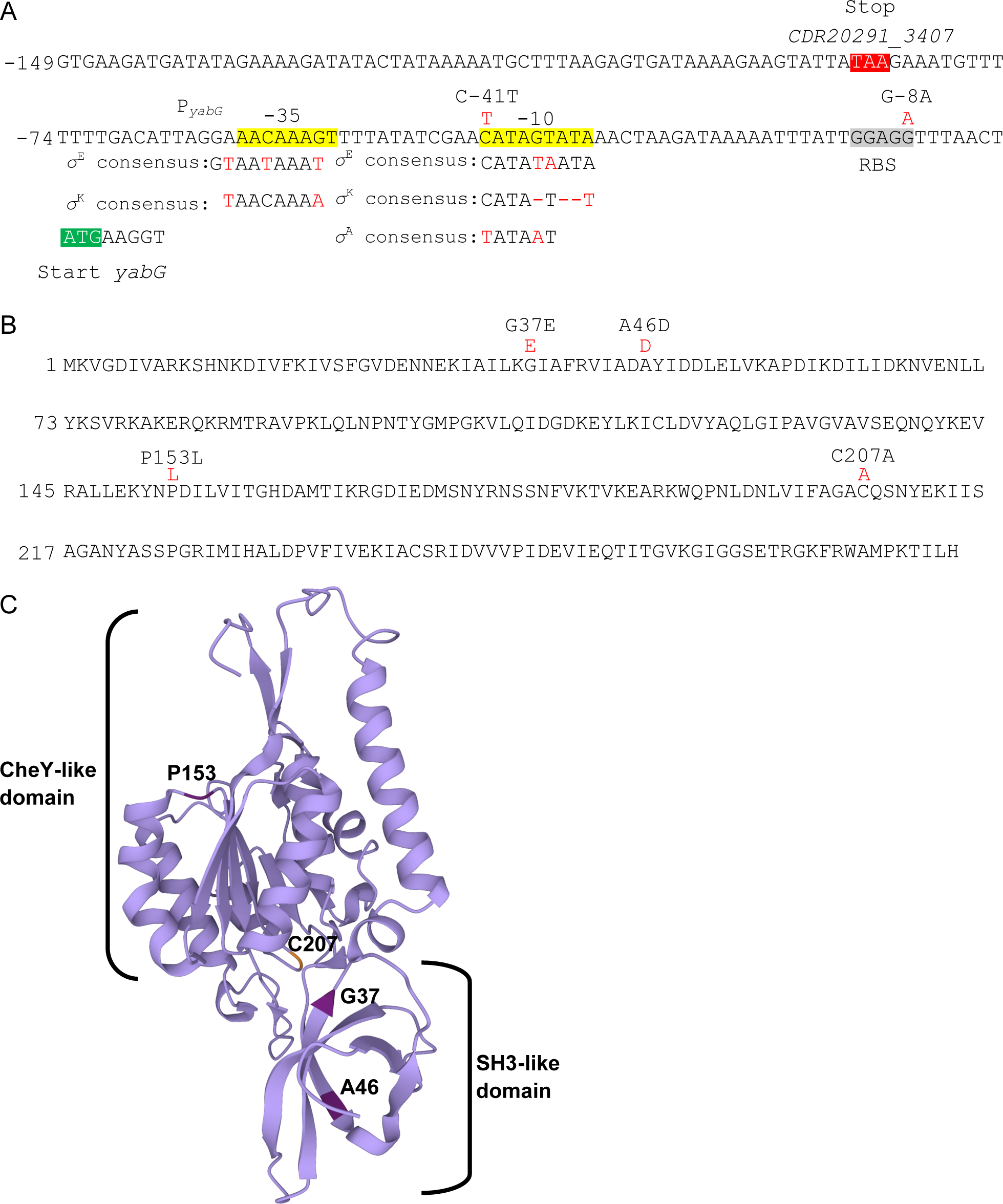
Mutations in the regulatory region of *yabG*. A) The −10 and −35 regions of the *yabG* promoter, utilized by both σ^E^ and σ^K^ in *C. difficile*, are shown in yellow and the consensus for the two σ factors indicated below. The position of the C-41T and G-8A mutations is shown in red; the C-41T substitution makes the −35 region of the *yabG* promoter closer to the consensus for σ^A^ recognition (shown below the sequence). The sequence shown corresponds to the fragment fused to the SNAP^Cd^ reporter with numbers indicating the position relative to the *yabG* start codon. B) The protein coding sequence of *C. difficile* YabG, with mutations, *yabG*_G37E_, *yabG*_A46D_, *yabG*_P153L_, and *yabG*_C207A_ shown in red. Numbers indicate the position relative to the fMet. C) Alphafold2 prediction of *C. difficile* YabG. G37, A46, and P153 are colored dark purple. The catalytic residue, C207, is colored in orange.

Here, we quantified the processing abilities of recombinantly expressed *yabG* mutant alleles and determined the *in vivo* effects on *C. difficile* spore germination. We show that of the previously identified *yabG* alleles, only YabG_A46D_ showed any activity in processing preproSleC. Mutations in the *yabG* locus result in misprocessing of germination proteins and spore coat proteins, whereas mutations in the region upstream of the *yabG* start codon (promotor and Shine-Dalgarno) affected the timing of expression.

## Results

### Expression of *YabG_A46D_* leads to reduced processing of preproSleC

YabG cleaves the pre-peptide from preproSleC, and proSleC is incorporated into the mature spore (Shrestha *et al*., 2019, Kevorkian *et al*., 2016, Marini *et al*., 2023). To understand the processing capabilities of the *yabG* EMS mutant alleles previously identified, we recombinantly expressed and purified *C. difficile* preproSleC in *E. coli*. preproSleC was incubated with *E. coli* lysate expressing YabG, YabG_C207A_, YabG_A46D_, YabG_P153L_, or YabG_G37E_ from an IPTG-inducible promoter. Samples were taken and protein separated by SDS-PAGE. The amount of proSleC was determined by immunoblotting and quantified by Li-COR imaging. Lysate expressing wild type YabG efficiently processed preproSleC to proSleC to nearly 100% within 1 hour of incubation (Figure 2A and S1A). YabG_C207A_, a catalytically inactive mutant, served as a negative control (Figure 2B, S1B) (Marini *et al*., 2023). Interestingly, of the mutant alleles, only YabG_A46D_ showed any ability to process preproSleC to proSleC. However, only 50% proSleC was present after 3 hours of incubation, suggesting that YabG_A46D_ does not process preproSleC as efficiently as the wild type allele (Figure 2C and S1C). YabG_G37E_ (Figure 2D and S1D) and YabG_P153L_ (Figure 2E and S1E) did not process preproSleC. During our analyses, we noticed the presence of a proSleC even in the absence of exposure to *E. coli* lysate (Figure S1, S2A and S2C). To ensure that the presence of proSleC in these assays was a product of preproSleC purification and not due to processing during incubation with *E. coli* lysate, lysate of *E. coli* BL21(DE3) containing an empty pET22b vector (EV) was incubated with the purified preproSleC. Processing was compared to the wild type *yabG* allele control (Figure S2B and S2D). Because the abundance of the proSleC form does not increase in the *E. coli* BL21(DE3) pEV control, we conclude that *E. coli* is not processing preproSleC and that the observed proSleC is a degradation product generated during our preproSleC purification. The presence of this product does not change the outcome or conclusions of these assays.

**Figure 2:**
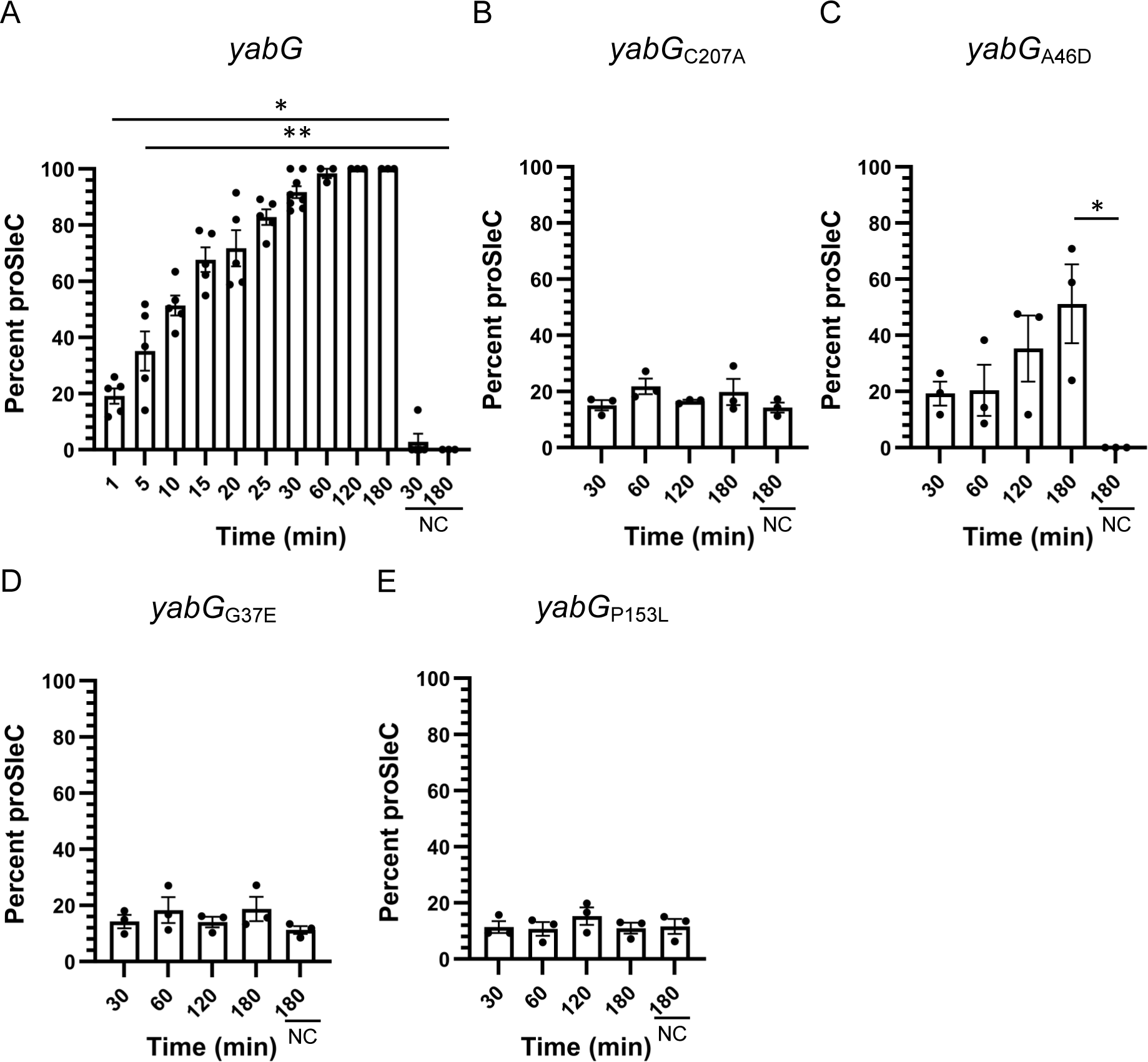
Processing of preproSleC by mutant YabG variants. Percent of proSleC present when incubated with the different *yabG* allele. The percentage of proSleC processed by: A) Wild type *yabG,* B) *yabG*_C207A_, C) *yabG*_A46D_, D) *yabG*_G37E_ and E) *yabG*_P153L_, were quantified on the LI-COR Odyssey-CLx, (proSleC signal)/(proSleC signal + preproSleC signal) *100. (C) indicates where no *E. coli* lysate was added. The data represent the averages from three to eight biological replicates and error bars represent the standard error of the mean. Statistical significance was determined using one-way ANOVA with Šídák’s multiple comparisons test (* p < 0.0332; ** < 0.0001). *Note: the purified SleC used in the assays with *yabG*_C207A_, *yabG*_G37E_, and *yabG*_P153L_ had ∼15% proSleC present prior to the assay in the sample as seen in the purified preproSleC negative controls (NC).

### YabG can shift the processing site of preproSleC to nearby arginine residues

We previously identified the YabG processing site of preproSleC to be immediately after R119 using Edman degradation (Shrestha *et al*., 2019). Interestingly, the deletion of the identified SRQS sequence resulted in processing of preproSleC after R115 and normal incorporation of proSleC into the spores. This suggested that YabG could shift its processing site to a nearby arginine (Shrestha *et al*., 2019). To test the limits of YabG to shift its processing site, we first co-incubated preproSleC_R119A_ with YabG, as described above. Because the preproSleC_ΔSRQS_ mutant was processed at R115 instead of R119, we hypothesized that the R119A protein would also be processed at R115 (Shrestha *et al*., 2019). Indeed, the first 5 amino acids of the resulting proSleC were SFSAQ, suggesting that YabG processed after R115 (Figure 3A). Next, we generated a mutant allele with both arginine amino acids replaced, preproSleC_R119A/R115A_, which YabG processed at R112 (Figure 3A). When a triple substitution was tested, preproSleC_R119A/R115A/R112A_, we observed no processing (Figure 3A). This suggests that YabG processes after arginine residues and that, upon mutation, the processing site shifts within certain limits.

**Figure 3:**
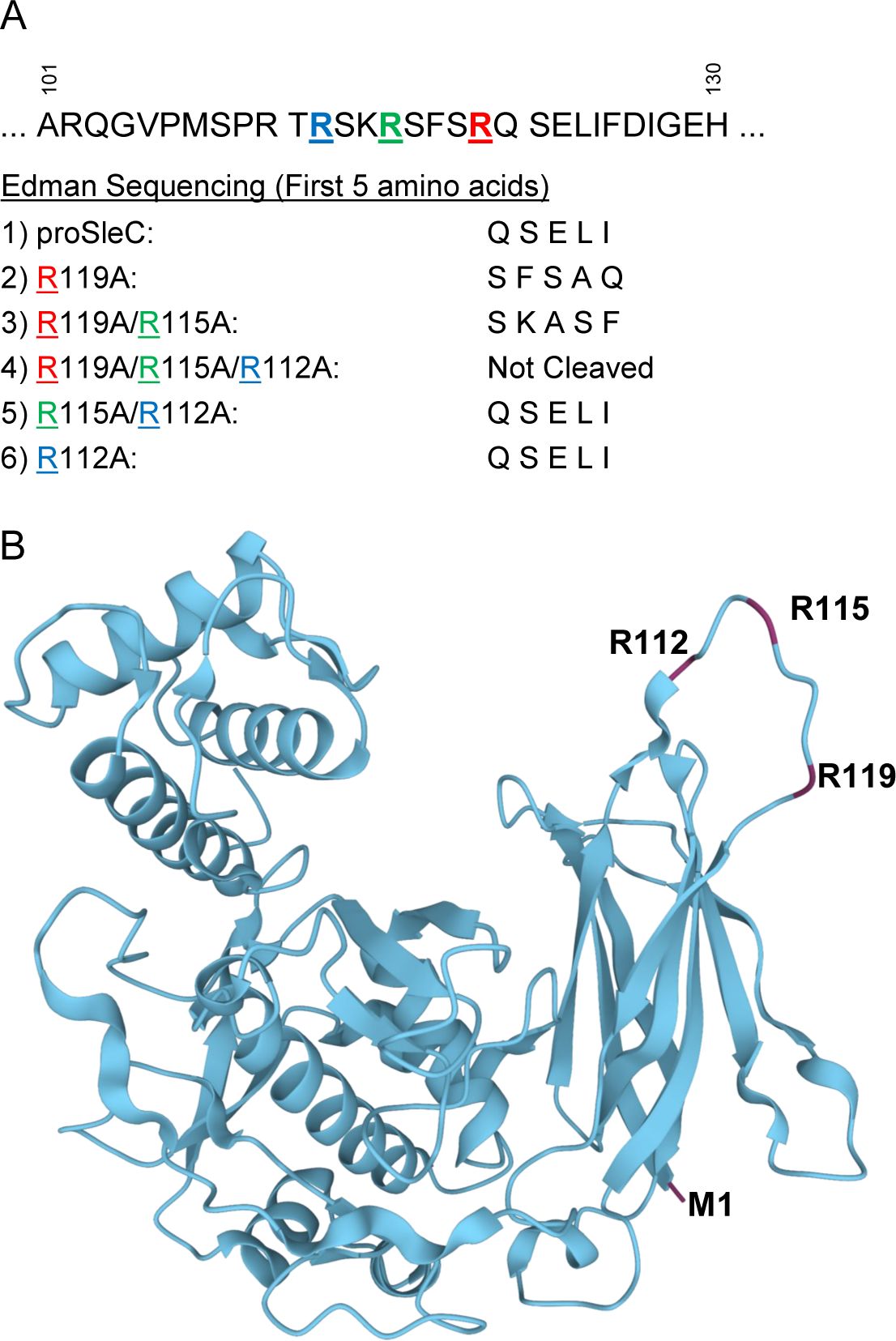
Processing site of recombinant preproSleC by *yabG* alleles. A) Recombinantly expressed and purified preproSleC, preproSleC_ΔSRQS_, preproSleC_R119A_, preproSleC_R119A/R115A_, preproSleC_R119A/R115A/R112A_, preproSleC_R115A/R112A_, and preproSleC_R112A_were incubated with *E. coli* lysate expressing YabG. The N-terminus of the YabG-cleaved preproSleC (proSleC) was determined by Edman sequencing. Arginine after which YabG processes preproSleC are colored at position 119 (red), 115 (green), and 112 (blue). B) *C. difficile* 630Δ*erm* SleC AlphaFold2 protein structure prediction. Highlighted are R112, R115, and R119 at which YabG cleaves and M1 (purple).

Because Edman degradation only reveals N-terminal amino acids, it was unknown if YabG was processing at these arginine residues in a sequential manner, *i.e.* at R112, then R115, and lastly R119. To test this hypothesis, preproSleC_R112A_ and preproSleC_R112A/R115A_ were recombinantly expressed, purified, and incubated with YabG. Edman degradation revealed that YabG processed both R115A and R115A/R112A alleles at R119 (Figure 3A). Using an AlphaFold2 prediction of the preproSleC protein structure, we noticed that the three arginine residues at which YabG processes are in a disordered region (Figure 3B) (Varadi *et al*., 2024, Jumper *et al*., 2021). Surprisingly, there is another arginine, R110, within this region that was not cleaved by the in any of our assays. Some AlphaFold2 models predict that the R110 is located within a beta sheet, suggesting that YabG only processes after arginine residues found in the disordered loops (Figure 3B). Alternatively, the presence of a proline at position 109 could be excluding YabG from processing at R110.

### Mutations in yabG result in a TA only germination phenotype

YabG processes two proteins involved in *C. difficile* spore germination: preproSleC and CspBA (Shrestha *et al*., 2019, Kevorkian *et al*., 2016). In prior work, we found that a *C. difficile yabG::ermB* mutant germinated in the presence of taurocholic acid (TA) alone and did not respond to the addition of a co-germinant (Shrestha *et al*., 2019). To determine if the mutations in *yabG* affected the ability to respond to co-germinants, isogenic mutants were made of *yabG*_A46D_, *yabG*_G37E_, *yabG*_P153L_, *yabG*_C207A_, *yabG*_C-41T_, and *yabG*_G-8A_ in a *C. difficile* R20291 Δ*pyrE* strain. Spores derived from the indicated strains in buffer alone did not germinate (Figure 4A and 4D). Wild type *C. difficile* Δ*pyrE* does not germinate with the addition of TA alone (Figure 4B and 4E) and requires a co-germinant (glycine) to germinate (Figure 4C and 4F). However, spores derived from the *C. difficile* Δ*pyrE yabG*_C207A_, *C. difficile* Δ*pyrE yabG*_A46D_, *C. difficile* Δ*pyrE yabG*_P153L_, and *C. difficile* Δ*pyrE yabG*_G37E_ strains germinated with the addition of TA but no co-germinant (Figure 4B). Spores derived from the *C. difficile* Δ*pyrE yabG*_G-8A_ and *C. difficile* Δ*pyrE yabG*_C-41T_ mutant strains also germinated in the presence of TA alone (Figure 4E). Germination of all the *C. difficile yabG* mutant spores was not enhanced by the addition of glycine (Figure 4C and F). Together these results support the previous findings that these mutations in *yabG* result in spores that no longer respond to co-germinants for germination.

**Figure 4:**
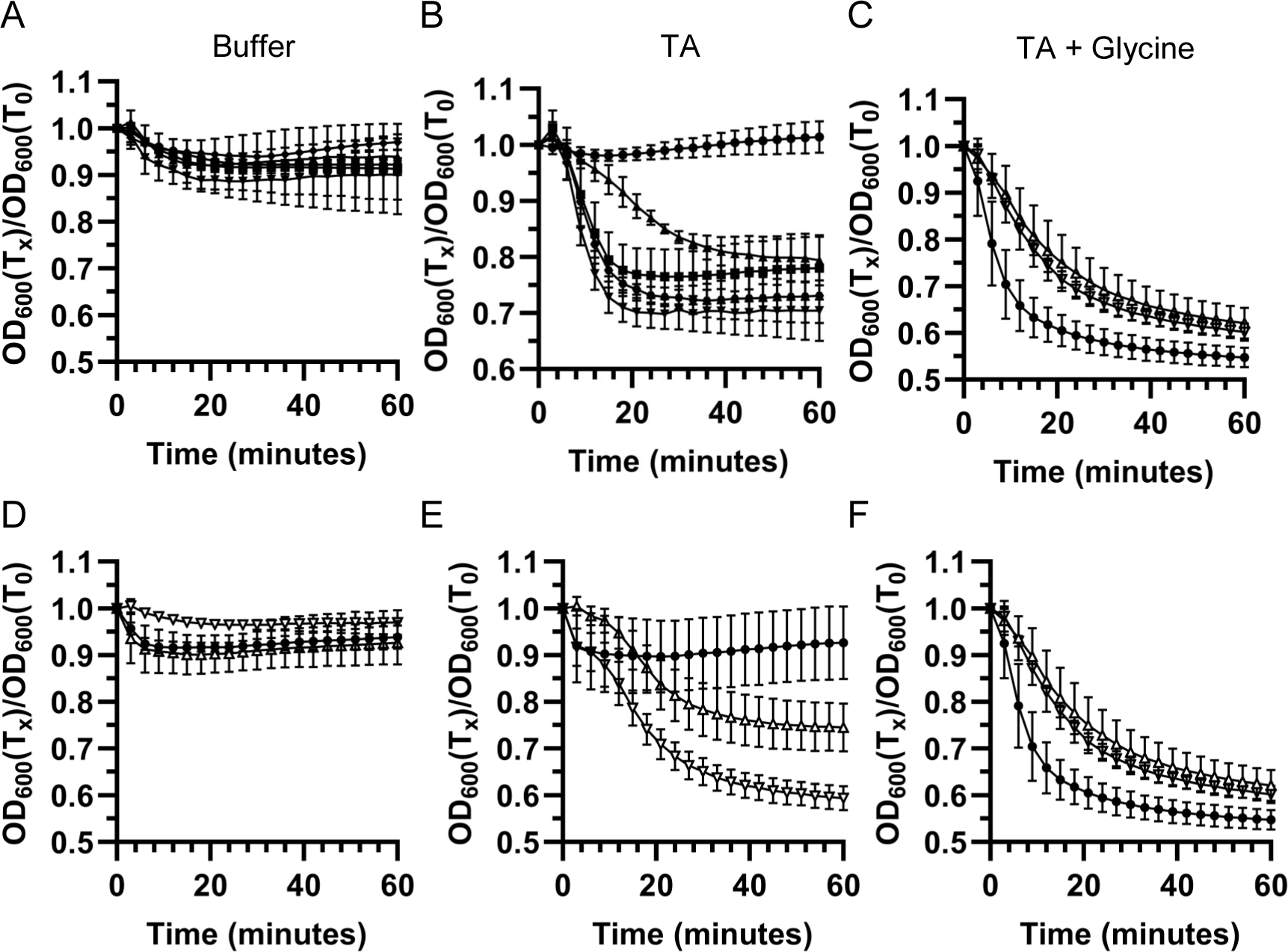
Mutations in YabG allow for germination in the presence of taurocholic acid only. Spores derived from *C. difficile* Δ*pyrE*, *C. difficile* Δ*pyrE; yabG*_C207A_, *C. difficile* Δ*pyrE; yabG*_A46D_, *C. difficile* Δ*pyrE; yabG*_P153L_, *C. difficile* Δ*pyrE; yabG*_G37E_, *C. difficile* Δ*pyrE; yabG*_C-8T_, and *C. difficile* Δ*pyrE; yabG*_G-41A_ germinate in the presence of taurocholic acid alone and do not require a co-germinant. 5 µL of OD_600_ = 100 spores derived from *C. difficile* Δ*pyrE* (circles), *C. difficile* Δ*pyrE yabG*_C207A_ (triangle), *C. difficile* Δ*pyrE yabG*_A46D_ (square), *C. difficile* Δ*pyrE yabG*_P153L_ (diamond), *C. difficile* Δ*pyrE yabG*_G37E_ (inverted triangle), *C. difficile* Δ*pyrE; yabG*_C-8T_ (inverted open triangle), and *C. difficile* Δ*pyrE; yabG*_G-41A_ (open triangle) were added to 95 µL germination buffer. A and D) Strains were suspended in buffer alone, B and E) in buffer supplemented with 10 mM TA, or C and F) in buffer supplemented with 10 mM TA and 30 mM glycine. Germination was monitored at OD_600_. Data points represent the averages from biological triplicate experiments and error bars represent the standard error of the mean.

### Mutations in yabG affect the abundance of YabG processed proteins

YabG is required for the expression of CotA and CdeM, a coat and an exosporium protein, respectively (Marini *et al*., 2023). In a *C. difficile yabG* mutant, these proteins have reduced abundance, and the spore has morphological changes (Marini *et al*., 2023). To investigate if our *C. difficile yabG* mutant alleles also exhibit changes to cortex/coat/exosporium protein abundance, we performed western blots on spore extracts. YabG was detected in the spore extracts of *C. difficile* Δ*pyrE yabG*_G37E_, *C. difficile* Δ*pyrE yabG*_A46D_, *C. difficile* Δ*pyrE yabG*_P153L_, and *C. difficile* Δ*pyrE yabG*_C207A_. *C. difficile* Δ*pyrE yabG*_C207A_ spores exhibited the highest amount of YabG, consistent with previous findings that it does not autoprocess (Figure 5A) (Marini *et al*., 2023). Of the three single amino acid substitution mutants, *C. difficile* Δ*pyrE yabG*_P153L_ showed the highest abundance of YabG, followed by *C. difficile* Δ*pyrE yabG*_G37E_, and *C. difficile* Δ*pyrE yabG*_A46D_ (Figure 5A). *C. difficile* Δ*pyrE yabG*_P153L_ and *C. difficile* Δ*pyrE yabG*_G37E_ had an abundant processed proSleC, differing from *in vitro* assays. YabG was not detected in spore extracts derived from the wildtype, *C. difficile* Δ*pyrE yabG*_G-8A_, and *C. difficile* Δ*pyrE yabG*_C-41T_ mutant strains. This likely results from YabG auto-proteolytic activity combined with reduced expression in the *yabG*_G-8A_ and *yabG*_C-41T_. Compared to wildtype, we also detected a greater abundance of the preproSleC form in all mutants tested (Figure 5A).

**Figure 5.**
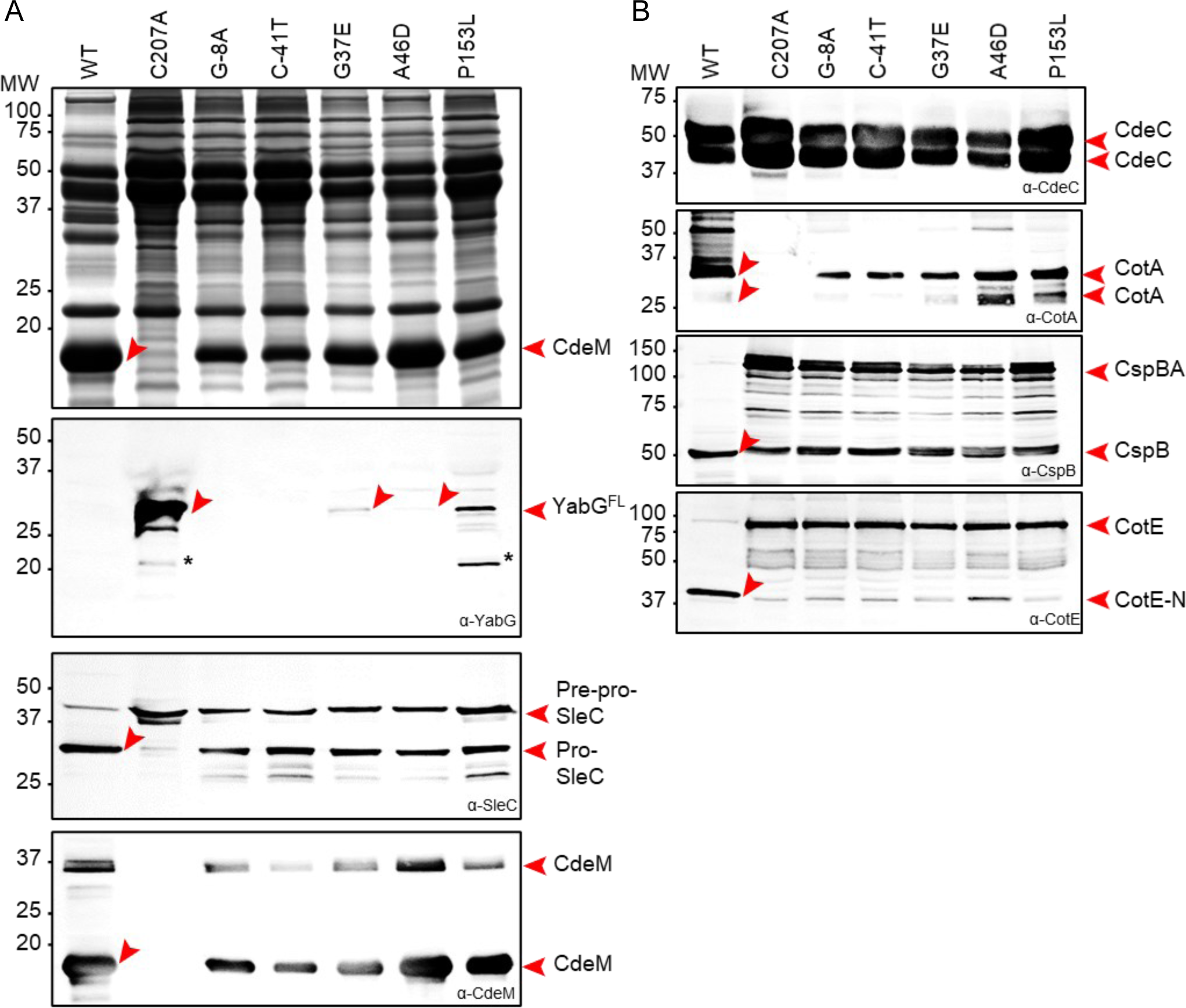
*yabG* mutations affect the level and/or extractability of coat and exosporium proteins. Spores of the WT, *yabG*_C207A_, and the various *yabG* point mutants, as shown, were purified, fractionated and the coat/exosporium and cortex/core proteins extracted. The proteins were resolved by 15% SDS-PAGE and the gels stained with Coomassie or subjected to immunoblot analysis with anti-YabG, anti-SleC, anti-CdeM (A), anti-CdeC, anti-CotA,anti-CspBA and anti-CotE antibodies (B). The position of the proteins and their main forms, recognized by the different antibodies is indicated by the red arrowheads. The position of molecular weight (MW) markers (in kDa) is indicated on the left side of the panel.

CdeM and CotA were absent from spores derived from the *yabG*_C207A_ strain (Figure 5A and 5B). We also observed less CdeM in spores of *C. difficile* Δ*pyrE yabG*_C-41T_, *C. difficile* Δ*pyrE yabG*_G-8A_, and *C. difficile* Δ*pyrE yabG*_G37E_ (Figure 5A). There was no effect on the observed CdeC (Figure 5B). The abundance of cleaved CotE was also reduced in all mutant strains. The *C. difficile* Δ*pyrE yabG*_A46D_ strain had increased processing compared to the other mutant strains. This is consistent with this allele retaining some activity against preproSleC (Figure 2C) (Shrestha *et al*., 2019). Finally, the autoprocessed form of CspB was also detected in all of the mutant spore extracts, consistent with previous findings in spores derived from a *C. difficile* Δ*yabG* mutant strain (Figure 5B) (Kevorkian *et al*., 2016).

### Mutations in yabG affect the spore morphology

To understand if spore assembly was affected in the mutant strains, we imaged the spores by transmission electron microscopy (TEM). In spores derived from the *C. difficile yabG*_C207A_ mutant, a loose connection between the cortex and coat layers is occasionally seen (Figure S3) (Marini *et al*., 2023). Also, the spore surface lacks electron density or has a very thin electron dense exosporium which lacks bumps; the appendage, when present, is not electron dense but rather shows a lamellar structure (Figure S3). In *C. difficile* Δ*pyrE yabG*_P153L_, *C. difficile* Δ*pyrE yabG*_A46D_ and *C. difficile* Δ*pyrE yabG*_G37E_ spores, the exosporium was thin, and, when present, the bumps were much smaller than in the wild type; the appendage region was less electron dense than in the wild type and showed a lamellar pattern (Figure S3). Spores of the *C. difficile* Δ*pyrE yabG*_G-8A_ and *C. difficile* Δ*pyrE yabG*_C-41T_ strains appeared similar. These observations are consistent with the idea that construction of the electron dense outer layer of the spore body and appendage is dependent on CdeM (Marini *et al*., 2023). In the strains which we observed a reduction in the electron density of the exosporium, we also observed a lower abundance of CdeM in the spore (Figure 5A), supporting this hypothesis. In all the spores, filamentous projections are seen emanating from the spore surface, which may correspond to the structures formed by the Bcl proteins (Pizarro-Guajardo *et al*., 2014, Paredes-Sabja *et al*., 2014, Pizarro-Guajardo *et al*., 2020).

### Characterizing the effects of yabG promoter mutations on yabG expression

To study the timing of *yabG* expression during sporulation, the fragment derived from the *C. difficile yabG* regulatory region (Figure 1A) was fused to the SNAP^Cd^ reporter. In the SNAP^Cd^ constructs, the *C. difficile yabG* RBS was fused to the start codon of the reporter so that mutations presumably affecting transcription (*C. difficile* Δ*pyrE yabG*_C-41T_) or translation (*C. difficile* Δ*pyrE yabG*_G-8A_) could be assessed (Figure 6A). These experiments were conducted in *C. difficile* 630Δ*erm* due to better data on the timing of *C. difficile* sporulation in this strain (Marini *et al*., 2023). As observed previously, the wild type promoter is active during late stages of sporulation, mainly in sporangia of phase grey/phase bright spores (Figure 6A, white arrows) (Marini *et al*., 2023). The *C. difficile* Δ*pyrE yabG*_C-41T_ mutation increases expression of the reporter earlier than the wild type (when the forespore is not yet discernible by phase contrast microscopy) (Figure 6A, black/grey arrows) and expression is reduced in sporangia at late stages in sporulation (Figure. 6A, white arrows). In contrast, in the *C. difficile* Δ*pyrE yabG*_G-8A_ mutant, production of the reporter had undetectable levels of expression at all stages of sporulation (Figure 6A). We next quantified the level of expression in sporangia of phase grey/phase bright spores of wild type *C. difficile* Δ*pyrE*, *C. difficile* Δ*pyrE yabG*_G-8A_, and *C. difficile* Δ*pyrE yabG*_C-41T_ strains at various stages (early, phase dark, phase bright) of sporulation (Figure 6B). Consistent with observations in Figure 6A, we observed *yabG*_C-41T_ expression earlier than other strains. *yabG*_C-41T_ expression was lower in abundance at later stages of sporulation (Figure 6B) compared to wild type, with the *yabG_G-8A_* allele showing the lowest expression in all stages of sporulation (Figure 6B).

**Figure 6.**
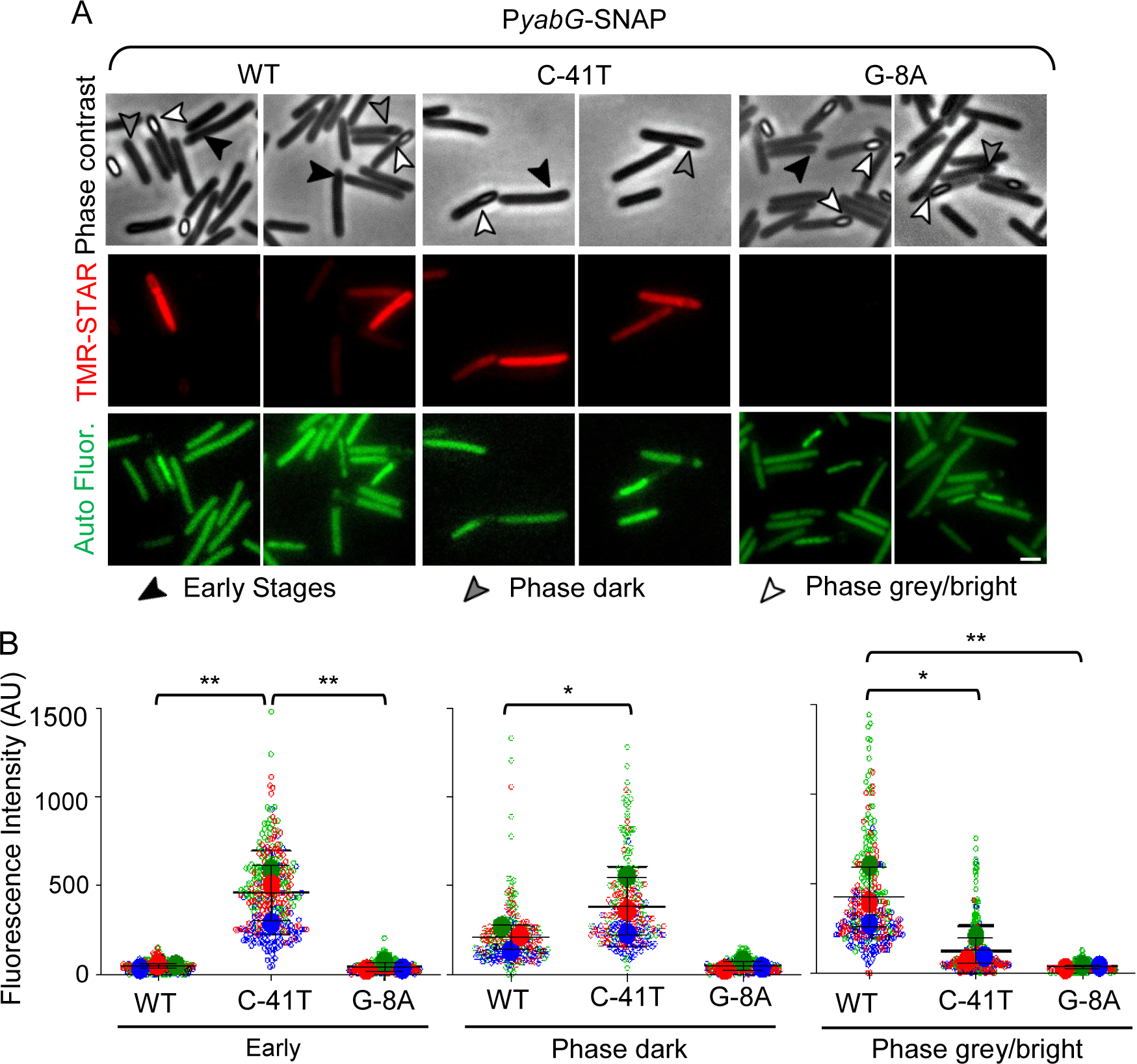
Sing cell analysis of P*yabG* expression. A) *C. difficile* sporulating cells carrying fusions of the WT *yabG* promoter or alleles with the G-8A or C-41T mutations fused to the *SNAP^Cd^*reporter in an otherwise WT background (630Δ*erm*) were collected after 20 hours of growth on 70:30 agar plates. The cells were stained with the SNAP substrate TMR-Star and examined by phase contrast (top panels) and fluorescence microscopy (middle panel, SNAP^Cd^ signal; bottom panels, autofluorescence). The black arrowheads show cells at early stages of sporulation, with no visible signs of a forespore; the grey arrowheads show sporangia of phase dark forespores and the white arrowheads indicate sporangia of phase grey or phase bright forespores. Scale bar, 1 μm. B) Intensity of the fluorescence signal per cell for the P*_yabG_*-*SNAP^Cd^* fusions described in A) in cells at early stages of sporulation or in sporangia of phase dark or phase grey/phase bright forespores. Fluorescence intensity is shown in arbitrary units (AU). The data from three independent experiments was represented using SuperPlots, with each color corresponding to a replicate (Lord *et al*., 2020). Statistical analysis was carried out using the ordinary one-way ANOVA and Tukey’s test. Asterisks correspond to p-values of p<0.05 (*) or p<0.01 (**).

## Discussion

Previous work on YabG has focused mostly on *Bacillus subtilis* (Takamatsu *et al*., 2000a, Takamatsu *et al*., 2000b, Kuwana *et al*., 2006, Yamazawa *et al*., 2022). Like in *C. difficile*, *B. subtilis* YabG is a sporulation-specific cysteine protease involved in the processing of several spore proteins during sporulation. Most target proteins of *B. subtilis* YabG (CotF, CotT, YeeK, YxeE, and SafA) are not conserved in *C. difficile* (Takamatsu *et al*., 2000a, Takamatsu *et al*., 2000b, Sebaihia *et al*., 2006). Additionally, the mechanism of target recognition and any additional targets remain unknown in *C. difficile*.

In *C. difficile,* YabG processes CspBA to CspB and CspA and preproSleC to proSleC (Marini *et al*., 2023, Shrestha *et al*., 2019, Kevorkian *et al*., 2016). However, in spores derived from a *C. difficile yabG* mutant, some CspB is present (Kevorkian *et al*., 2016). This CspB form is due to either autoprocessing or another protease that is capable of processing CspBA (Kevorkian *et al*., 2016, Marini *et al*., 2023, Shrestha *et al*., 2019). A previous study found a decrease in the abundance of SpoIVA in spores derived from a *C. difficile yabG* mutant, but the involvement of YabG in the incorporation of SpoIVA into mature spores is poorly understood (Kevorkian *et al*., 2016). Because *C. difficile* YabG is the only protease with the ability to cleave preproSleC, and the preproSleC form is the only form incorporated into mature spores in *C. difficile yabG* mutants, preproSleC was used as our target to quantify the processing capabilities of YabG, YabG_C207A_, YabG_A46D_, YabG_P153L_, and YabG_G37E_ (Kevorkian *et al*., 2016, Marini *et al*., 2023, Shrestha *et al*., 2019). We found that YabG is very efficient and processed ∼50% of preproSleC to proSleC within 10 minutes and ∼100% within 1 hour of *in vitro* incubation (Figure 2A, S1B). However, YabG_A46D_ only processes 50% of preproSleC within 3 hours of *in vitro* incubation (Figure 2C, S1C). Interestingly, YabG_P153L_ and YabG_G37E_ did not show any activity at the times tested (Figure 2 and S1). Because only one target of YabG was tested, it is possible that these alleles are more active on other unidentified YabG targets. We consider this unlikely because of the processing observed in spores (Figure 5). CspBA is another YabG target, but some CspBA is processed and incorporated into mature spores of a *C. difficile* Δ*yabG* mutant strain. This suggests that another protease can cleave CspBA to CspB and CspA (Shrestha *et al*., 2019, Kevorkian *et al*., 2016). Spores derived from the *C. difficile* R20291 *yabG::ermB* mutant strain only contained CspBA and not CspB and CspA (Shrestha *et al*., 2019) but our isogenic mutants (Figure 5B) contain some processed CspBA suggesting differences between the two *C. difficile* strains (*C. difficile* R20291 vs. *C. difficile* R20291 Δ*pyrE*) or because *yabG* is not produced in the *C. difficile* R20291 *yabG::ermB* mutant where as it is produced in *C. difficile* R20291 Δ*pyrE yabG*_C207A_, but not catalytically active.

The processing site of YabG on preproSleC had previously been identified as R119. However, two other arginine residues are near R119, and we wanted to determine if YabG could also process at those sites (Shrestha *et al*., 2019). Interestingly, we found that upon mutation of preproSleC R119 to alanine, processing occurred at R115. In the preproSleC R119A/R115A mutant, processing occurred at R112. No processing was observed for the combined triple mutant, R119A/R115A/R112A. Interestingly, YabG does not cleave at R110 and in a predicted AlphaFold2 structure of preproSleC shows R119, R115, and R112 all reside within an exposed, unstructured loop, whereas R110 resides within a beta strand, mainly excluding it from processing (Figure 3B). It could also be the presence of P109 that excludes R110 from YabG processing.

All the spores derived from the *C. difficile* Δ*pyrE yabG* mutants germinated with TA only and did not require a co-germinant in order to germinate, unlike the parental *C. difficile* Δ*pyrE* strain (Figure 4). These results are consistent with previous findings. Interestingly, *C. difficile* Δ*pyrE yabG*_C207A_ appears to germinate more slowly than the other mutants in TA only conditions. However, this may be due to the complete loss of YabG catalytic activity, resulting in a lack of processing of alterative targets (Shrestha *et al*., 2019).

Spores derived from the *C. difficile* Δ*yabG* and *C. difficile yabG*_C207A_ strains incorporate less CotA and CdeM (Figure 5A and 5B) (Marini *et al*., 2023). CotA is a component of the spore coat and is required for assembly of the outer surface layers of the spore (Permpoonpattana *et al*., 2013, Permpoonpattana *et al*., 2011). CdeM is a component of the exosporium (Calderon-Romero *et al*., 2018). Upon investigation of *cdeC* and *cdeM* transcript levels of *C. difficile* Δ*yabG* sporulating cells, their levels were lower compared to wild type, suggesting YabG was regulating their transcription, though the mechanism by which this occurs remains unknown (Marini *et al*., 2023). Therefore, to investigate if the C. *difficile yabG* alleles also affect spore cortex/coat/exosporium proteins, western blots were performed against CdeM, CdeC, CotA, CotE, CspBA, preproSleC, and YabG. CdeM was less abundant or less extractable from the purified spores of *C. difficile* Δ*pyrE yabG*_C-41T_ (Figure 5A). In *C. difficile* Δ*pyrE yabG*_C-41T_ expression of the reporter is even higher than in the wild type (Figure 6B), which could potentially be due to earlier expression of *yabG* interfering with transcription of *cdeM* and *cotA* and/or assembly of the proteins (Figure 5). Interestingly, the levels/extractability of CdeM and CotA, were higher in *C. difficile* Δ*pyrE yabG*_A46D_ and *C. difficile* Δ*pyrE yabG*_P153L_ spores, as compared to *C. difficile* Δ*pyrE yabG*_G37E_ spores (Figure 5A and 5B). One possibility is that the *yabG*_A46D_ and *yabG*_P153L_ alleles are more efficient than the *yabG*_G37A_ allele at promoting *cdeM* and *cotA* transcription. Processing of preproSleC, CspBA, and CotE is incomplete (Figure 5A and 5B), and we do not know the location or timing of these events. Because proSleC and CspB and CspA are found in the cortex, we hypothesize that they are processed in the cytoplasm before being transported to the cortex (Baloh *et al*., 2022).

YabG was not detected in the promoter mutant *C. difficile* Δ*pyrE yabG*_C-41T_ or the RBS mutant *C. difficile* Δ*pyrE yabG*_G-8A_, similar to wild type. This is most likely due to its auto-proteolytic activity, however the lack of YabG in *C. difficile* Δ*pyrE yabG*_G-8A_ is likely due to loss of expression (Figure 5A and Figure 6B) (Marini *et al*., 2023). In all *yabG* mutants with a single amino acid substitution (G37A, A46D, and P153L), YabG was detected in the spore extracts, but at levels lower than the catalytically inactive YabG_C207A_ allele. This suggests that these variants are less active than the WT protein (Figure 5A). Consistent with the lower activity of YabG_P153L_, autoprocessing and processing of preproSleC, CspBA, and CotE was diminished relative to *C. difficile yabG*_G37E_ and *C. difficile yabG*_A46D_ (Figure 5A and 5B).

*C. difficile* Δ*pyrE yabG*_A46D_ showed the lowest amount of YabG in spore extracts of the three single amino acid substitution mutants. This is consistent with YabG_A46D_ showing activity against purified preproSleC *in vitro* (Figure 2C and 5A). Reduced activity of YabG can result from single amino acid substitutions in different parts of the protein (A46D and G37E in the SH3 domain; P153L in the CheY-like, catalytic domain). Together this suggests that the different alleles have various levels of catalytic activity.

The main morphological changes in *C. difficile* Δ*yabG* or *C. difficile yabG*_C207A_ spores include a loose connection between the cortex and coat layers, a thin outermost electron dense layer (thought to be part of the exosporium), and a lack of electron density in the appendage region, which reveals an underlying pattern of lamellae (Marini *et al*., 2023). The thinner outer layer and the lack of electron density of the spore appendage is thought to be caused by the absence of CdeM, as it is not expressed in *yabG* mutants. Accordingly, the electron density of the appendage is restored when *cdeM* is expressed from the *yabG*-independent *cotE* promoter (Marini *et al*., 2023). *C. difficile yabG* mutants, were originally studies in *C. difficile* 630Δ*erm*, however, we used *C. difficile* R20291 Δ*pyrE* as a parental strain (Marini *et al*., 2023). A striking difference between spores of the two strains is that the outermost electron dense spore layer is much thicker in R20291 spores and presents bumps that protrude from the entire surface of the spore (Figure S3, wild type spores). Apart from the *C. difficile yabG*_C207A_ mutant, there were no overt signs of an impaired cortex/coat connection in the point mutation strains, suggesting that a lower level of YabG activity is sufficient to prevent this phenotype (Figure S3).

It remains unknown how YabG recognizes its targets in any organism. In *B. subtilis*, YabG processes coat proteins, however only SpoIVA has an orthologue in *C. difficile (Kuwana et al., 2006, Takamatsu et al., 2000a, Takamatsu et al., 2000b, Yamazawa et al., 2022).* Moreover, despite universal conservation of the protein in spore forming bacteria, the absence of conserved substrates is intriguing. Future work needs to be done to identify YabG substrates and reveal any conserved features that may help to identify substrates in other non-model, spore forming organisms.

### Experimental Procedures

#### Bacterial growth conditions

All bacterial strains are listed in Table S2*. C. difficile* strains were grown in a Coy anaerobic chamber at 37 °C, 3-4% H_2_, 5% CO_2_, and balanced N_2_ on either brain heart infusion (BHI) medium (Difco) supplemented with 0.1% L-cysteine and brain heart infusion supplemented with 5 g / L yeast extract and 0.1% L-cysteine (BHIS) or 70:30 media as indicated. When necessary, media was supplemented with thiamphenicol (10 µg / mL), kanamycin (50 µg / mL), D-cycloserine (250 µg / mL), tetracycline (5 µg / mL), uracil (2 µg / mL), theophylline (2 g / L) or taurocholate (TA) (0.1%). Defined minimal media for *C. difficile* (CDMM) supplemented with 5-fluoroorotic acid (FOA; 2 mg / mL) and uracil (5 µg / mL) was used for the selection of Δ*pyrE* mutants. *E. coli* strains were grown at 37 °C on LB medium supplemented with chloramphenicol (20 µg / mL) and or ampicillin (100 µg / mL) for plasmid maintenance. *E. coli* BL21 (DE3) was grown in 2x tryptone yeast (2XTY), medium supplemented with chloramphenicol (20 µg / mL) and or ampicillin (100 µg / mL) for plasmid maintenance and used for recombinant protein expression.

#### Plasmid constructions

Plasmids pMS17, pMS43, pMS44, and pMS45 were made from pMTL-YN4 as follows. PMTL-YN4 was digested with <u>MluI</u> / XhoI and purified by gel electrophoresis, extracted, then assembled with their respective regions of homology via Gibson assembly (Gibson *et al*., 2009). Regions of pMS17 homology were amplified from *C. difficile* R20291 with primers 5’ yabG C207A pMTL-YN4, 3’ yabG C207A upstream, 5’ yabG C207A downstream, and 3’ yabG C207A pMTL-YN4. Region of pMS43 homology were amplified from *C. difficile* mutant 20C using primer pairs 5’ yabG A46D_up / 3’ yabG A46D_up, and 5’ yabG A46D_down / and 3’ yabG A46D_down. Regions of pMS44 were amplified from *C. difficile* mutant 30A using primer pairs 5’ yabG P153L_up / 3’ yabG P153L_up and 5’ yabG P153L_down / 3’ yabG P153L_down, respectively. Regions of pMS45 homology were amplified from *C. difficile* mutant 30C using primer pairs 5’ yabG G37E_up / 3’ yabG G37D_up and 5’ yabG G37E_down / 3’ yabG G37E_down. Plasmids pJB86 and pJB87 were made from pJB81 as follows. pJB81 was digested by NotI / XhoI and purified by gel electrophoresis, extracted, then assembled with their respective regions of homology via Gibson assembly. Regions of homology for pJB86 were amplified with primers 5’ prom_C-8T_up / 3’ prom_C-8T_up and 5’ prom_C-8T_down / 3’ prom_C-8T_down, while those for pJB87 were amplified with primers 5’ yabG_G-41A_up / 3’ yabG_G-41A_up and 5’ yabG_G-41A_down / 3’ yabG_G-41A_down. After Gibson assembly and transformation into *E. coli* DH5α (Hanahan, 1983), colonies were re-streaked and tested via PCR for correct assembly, followed by whole plasmid sequencing

Plasmids pMS48, pMS49, pMS50, pMS51, pMS59, and pMS60 were made from pET22b as follows. pET22b was digested with NdeI / XhoI and purified by gel electrophoresis, extracted, then assembled with their respective regions of homology via Gibson assembly. Regions of pMS48 homology were amplified from *C. difficile* R20291 using primer pairs 5’ pET22b_yabG_up / 3’ yabG A46D_up and 3’ yabG A46D_up / 3’ pET22b_yabG_down. Regions of pMS49 homology were amplified from *C. difficile* mutant 30A using primer pairs primer pairs 5’ pET22b_yabG_up and 3’ pET22b_yabG_down. Regions of pMS50 were amplified from *C. difficile* mutant 30C using primer pairs 5’ pET22b_yabG_up and 3’ pET22b_yabG_down. Regions of pMS51 were amplified from *C. difficile* MRS05 using primer pairs 5’ pET22b_yabG_up and 3’ pET22b_yabG_down. Regions of pMS59 were amplified from *C. difficile* R20291 using primer pairs 5’ pET22b preproSleC / 3’ pET22b R112A and 5’ pET22b R112A and 3’pET_SleC. Regions of pMS60 were amplified from *C. difficile* R20291 using primer pairs 5’ pET22b preproSleC / 3’ pET22b R112A/R115A and 5’ pET22b R112A/R115A / 3’pET_SleC. After Gibson assembly and transformation into *E. coli* DH5α, colonies were re-streaked and tested via PCR for correct assembly, followed by whole plasmid sequencing

Plasmids pAC57, pAC58, and pAC59 were made from pET22b as follows. pET22b was digested with NdeI / BamHI and purified by gel electrophoresis, extracted, then assembled with their respective regions of homology via Gibson assembly. Regions of pAC57 were amplified from *C. difficile* R20291 using primers SleC 119A_Fp / pet22b_yabg_SleC6his_Rp and pet22b_SleC_Fp / SleC R119A_Rp. Regions of pAC58 were amplified from *C. difficile* R20291 using primer pairs SleC 119115112A_Fp / pet22b_yabg_SleC6his_Rp and pet22b_SleC_Fp / SleC 119115112A_Rp. Regions of pAC59 were amplified from *C. difficile* R20291 using primer pairs SleC 119115A_Fp / pet22b_yabg_SleC6his_Rp and pet22b_SleC_Fp / SleC 119115A_Rp. After Gibson assembly and transformation into *E. coli* DH5α, colonies were re-streaked and tested via PCR for correct assembly, followed by whole plasmid sequencing using Oxford Nanopore Technology by Plasmidsaurus inc. (Eugene, OR).

#### ΔpyrE mediated allelic exchange

*C. difficile* strains MRS04, MRS05, MRS06, and MRS07 were made as previously described (Ng *et al*., 2013) from parental strain KNM05 (McAllister *et al*., 2017) with plasmids, pMS43, pMS17, pMS45, and pMS44 respectively. Briefly, transconjugants were passaged on BHIS supplemented with thiamphenicol, uracil, D-cycloserine, and kanamycin. Integrants were passaged onto CDMM + FOA + uracil. Colonies were passaged onto BHIS and BHIS supplemented with thiamphenicol to screen for plasmid loss. Loss of plasmid was confirmed by PCR and whole genome re-sequencing was performed to confirm genotype.

#### Theophylline mediated allelic exchange

*C. difficile* strains JNB25 and JNB26 were made as previously described (Brehm & Sorg, 2024) from parental strain KNM05 with plasmids pJB87 and pJB86, respectively. Briefly, the plasmids were introduced into KNM05 via *E. coli* HB101 pRK24 conjugation and confirmed by PCR. Transconjugants were plated on BHIS Tm plates, where plasmid integration was screened by colony size. Colonies that contained the integrated plasmid were isolated and plated on BHIS containing 2 g / L theophylline to encourage plasmid excision. Colonies were isolated and tested for their respective mutations by PCR amplification and sequencing, followed by whole genome sequencing of the completed strains. Whole genome sequencing was performed on strains and sent for Illumina sequencing at SeqCoast Genomics LLC (Portsmouth, NH) and raw sequence reads are uploaded to the Sequence Read Archive under BioProject ID: PRJNA1112411.

#### SNAP^Cd^ fusions

To construct the P*yabG* transcriptional fusions to the SNAP^Cd^ reporter, the promoter regions of *yabG* were PCR-amplified using as the template genomic DNA of *C. difficile* R20291 Δ*pyrE* 31D (strain 4049030, bearing the C-8T mutation), *C. difficile* R20291 Δ*pyrE* 27E (strain 4049063, with the G-41A mutation) or *C. difficile* R20291Δ*pyrE* (lab strain AHCD774, *yabG*^WT^). The primer pairs used were PyabG-EcoRI-Fw and C8T-SNAP-SOE-Rev for the first strain and PyabG-EcoRI-Fw and YabG-978-SOE-Rev for the last two strains. The PCR reactions produced fragments of 271 bp. The SNAP^Cd^ gene was PCR amplified from pFT47 (Pereira *et al*., 2013) using primers SNAP-Fw and SNAPCd-*HindIII*-Rev to produce a fragment with 558 bp. The two fragments were joined by PCR using primers PyabG-EcoRI-Fw and SNAPCd-*HindIII*-Rev. This produced fragments of 809 bp which were cleaved using EcoRI and HindIII and inserted between the same sites of pMTL84121. This originated plasmids pCO39 (P*yabG*-SNAP^Cd^), pCO40 (P*yabG*^C8T^-SNAP^Cd^) and pCO41 (P*yabG*^G42A^-SNAP^Cd^). These plasmids were transformed into *E. coli* HB101 (RP4) originating strains AHEC1588 (pCO39), AHEC1589(pCO40) and AHEC1590(pCO41) and then transferred to *C. difficile* 630Δ*erm pyrE*^+^ (AHCD1190) by conjugation to produce strains AHCD1842, AHCD1843 and AHCD1844, respectively. All primers are listed in table S1. All strains and plasmids are listed in Table S2.

#### Germination assay

Germination was monitored using a Spectramax M3 plate reader (Molecular Devices, Sunnyvale, CA) (Shrestha *et al*., 2017, Bhattacharjee & Sorg, 2018, Shrestha & Sorg, 2018, Shrestha *et al*., 2019, Shrestha & Sorg, 2019). OD_600_ = 100 spores were heat activated for 30 min at 65 °C. 5 µL of spores was added to 95 µL of germination buffer containing 50 mM HEPES alone, or HEPES supplemented with 10 mM TA or with 10 mM TA and 30 mM glycine in a 100 µL total volume. The OD_600_ was monitored for 1 hour at 37 °C.

#### SleC expression and purification

Plasmid pKS08 was transformed into *E. coli Rosetta* BL21 (DE3) and incubated overnight at 37 °C on LB-agar supplemented with chloramphenicol and ampicillin. The plate was scraped into 1 mL of LB and used to inoculate 1 L of 2XTY supplemented with chloramphenicol and ampicillin in baffled flasks (OD_600_ = 0.01). The culture was incubated at 37 °C at 190 rpm until the OD_600_ was between 0.6 and 0.8, at which point the culture was induced with 250 µM IPTG and incubated for 16 hours at 16 °C and 130 rpm. Cultures were pelleted at 6,370 x g for 15 min at 4 °C. Supernatant was discarded and pellets were stored at −80 °C until use. 1 liter of cells was resuspended in 25 mL of 300 mM NaCl, 50 mM Tris-HCl, 15 mM imidazole (pH 7.5) and 0.03 mM PMSF. Each 25 mL of cells was supplemented with lysozyme and DNase I and was rocked for 30 minutes at 4 °C prior to sonication on ice at 27% amplitude for 20 min. Samples were then centrifuged at 25,900 x g for 30 min at 4 °C and the supernatant was combined with 1 mL of Ni-NTA Agarose beads. Samples were rocked overnight at 4 °C. Beads were washed twice with 300 mM NaCl, 50 mM Tris-HCl, 30 mM imidazole (pH 7.5). Beads were washed once with 300 mM NaCl, 50 mM Tris-HCl, 15 mM imidazole (pH 7.5) and then eluted with the same buffer but supplemented with 500 mM imidazole. Samples were concentrated using a 10 kDA molecular weight cut off centrifugal device. The protein was further purified by size exclusion chromatography using a Cytiva ÄKTA Pure system (Marlborough, MA) on a Superdex 200 increase 10/300 GL column and concentrated again. Protein was diluted to 2.5 mg / mL, aliquoted, flash frozen in a dry ice-ethanol bath, and stored at −80 °C until use.

#### YabG in vitro lysate preparation

Plasmids containing wild type (pAC28), mutant (pMS48, pMS49, pMS50, and pMS51) *yabG*, or an empty pET22b vector were transformed into *E. coli Rosetta* BL21 (DE3), and incubated overnight at 37 °C on LB-agar supplemented with chloramphenicol and ampicillin. The plate was scraped into 1 mL of LB and used to inoculate 50 mL of LB supplemented with chloramphenicol and ampicillin in 250 mL flasks, so the starting culture OD_600_ was at 0.01. The culture was incubated at 37 °C at 170 rpm until the OD_600_ was between 0.6 and 0.7. The culture was induced with 250 µM IPTG and incubated for 1 hour at 37 °C. YabG cultures were pelleted by centrifugation at 4,415 x g, for 15 min at 4 °C. The supernatant was discarded, and cells were resuspended in 4 mL of 300 mM NaCl, 50 mM Tris-HCl (pH 7.5) before sonication on ice at 27% amplitude for 20 min. Bacterial lysate was immediately used in the SleC processing assay.

#### SleC processing assay

Recombinantly expressed and purified preproSleC from *E. coli* was incubated with the indicated recombinantly expressed *yabG* allele lysate. Incubations were conducted at 37 °C for 1 min, 5 min, 10 min, 15 min, 20 min, 25 min, 30 min, 1 hour, 2 hours, or 3 hours. The reactions occurred in 25 µL total volume [56 µg preproSleC present in 22.5 µL + 2.5 µL of the indicated *yabG* allele or buffer 300 mM NaCl, 50 mM Tris-HCl, (pH 7.5)]. As a control, purified preproSleC was added to buffer alone. Samples were denatured at 95 °C for 20 minutes, diluted with 2x Nu-PAGE sample buffer, and stored at −20 °C until use.

#### Western blotting

340 ng or 34 ng of purified SleC from the SleC processing assays were separated by 12% SDS-PAGE. Protein was then transferred onto low-fluorescence polyvinylidene difluoride membrane (PVDF) using the BIO-RAD Trans-Blot Turbo Transfer system at 25 V, 1.0 A for 30 min. The membranes were blocked in 5% skim milk in Tris-buffered saline with 0.1% Tween-20 (TBST) rocking overnight at 4 °C. The membranes were incubated with anti-SleC antibody for 1 hour in 5% skim milk in TBST and washed three times with TBST. For the secondary antibody, LI-COR IRDye 680RD goat anti-rabbit antibody was used to label the membranes for 1 hour in 5% skim milk in TBST in the dark. The membranes were washed three times, in the dark, with TBST. Blots were imaged wet on the Odyssey® CLx and the Odyssey M imaging systems (Li-COR, Lincoln, NE) using the 700-800 nm channels. The fluorescent bands were visualized and analyzed using ImageStudio software version 5.2 or LI-COR Acquisition software version 1.2 and analyzed with Empiria Studio version 2.3 (LI-COR Biosciences).

#### Edman sequencing

Plasmids pKS08, pAC57, pAC59, pAC58, pMS60, and pMS59 were induced and protein was purified the same as described above. Purified preproSleC, preproSleC_R119A_, preproSleC_R119A/R115A_, preproSleC_R119A/R115A/R112A_, preproSleC_R115A/R112A_, and preproSleC_R112A_ were incubated with *E. coli* lysate expressing wild type *yabG* for three hours. The preproSleC processing assays were separated on a 12% SDS-PAGE and transferred to PVDF membrane using the BIO-RAD Trans-Blot Turbo Transfer system at 25 V, 1.0 A for 30 min. The membrane was thoroughly washed under dH_2_O for ∼ 10 minutes to remove residual glycine. The membrane was stained with Coomassie Brilliant Blue for 20 minutes (50% MeOH, 10% acetic acid, 40% dH_2_O, and 0.05% Coomassie). The membrane was destained (50% MeOH, 10% acetic acid, and 40% dH_2_O). The proSleC band was excised from the membrane using a fresh razor blade. The band was sent for Edman sequencing of the first five amino acids at The Protein Facility at Iowa State University.

#### SNAP^Cd^ labelling, phase contrast, fluorescence microscopy and image analysis

For SNAP labelling, cells were withdrawn from sporulating cultures after 20 hours of growth on 70:30 medium (Putnam *et al*., 2013); the samples were mixed for 30 min in the dark with the TMR-Star substrate (New England Biolabs) at a final concentration of 250 nM (Pereira *et al*., 2013). Cells were collected by centrifugation (4,000 x g for 5 min at room temperature), washed four times with 1 mL of phosphate-buffered saline (PBS; 137 mM NaCl, 10 mM Phosphate, 2.7 mM KCl, pH 7.4), and resuspended in 0.5 mL of PBS. For phase contrast and fluorescence microscopy, cells were mounted on 1.7% agarose coated glass slides and observed on a Leica DM6000B microscope equipped with a phase contrast Uplan F1 100x objective and captured with a CCD Andor Ixon camera (Andor Technologies). Images were acquired and analyzed using the Metamorph software suite (version 5.8; Universal Imaging) and adjusted and cropped using *ImageJ*.

#### Statistical analysis

All processing assays and germination assays were performed in at least biological triplicate and the data represents the averages from the data sets. Error bars represent the standard error of the mean. A one-way ANOVA Šídák’s multiple comparisons test was used to compare the quantified protein amounts.

## Acknowledgements

We thank members of the Sorg laboratory for critical comments during the preparation of this manuscript. We also thank Joel Nott at The Protein Facility of Iowa State University for performing the Edman sequencing.

## Funding and additional information

This project was supported by awards 5R01AI116895 and 5R01AI172043 to J.A.S. from the National Institute of Allergy and Infectious Diseases. The content is solely the responsibility of the authors and does not necessarily represent the official views of the NIAID. The funders had no role in study design, data collection and interpretation, or the decision to submit the work for publication.

---

**Supporting Figure 1:**
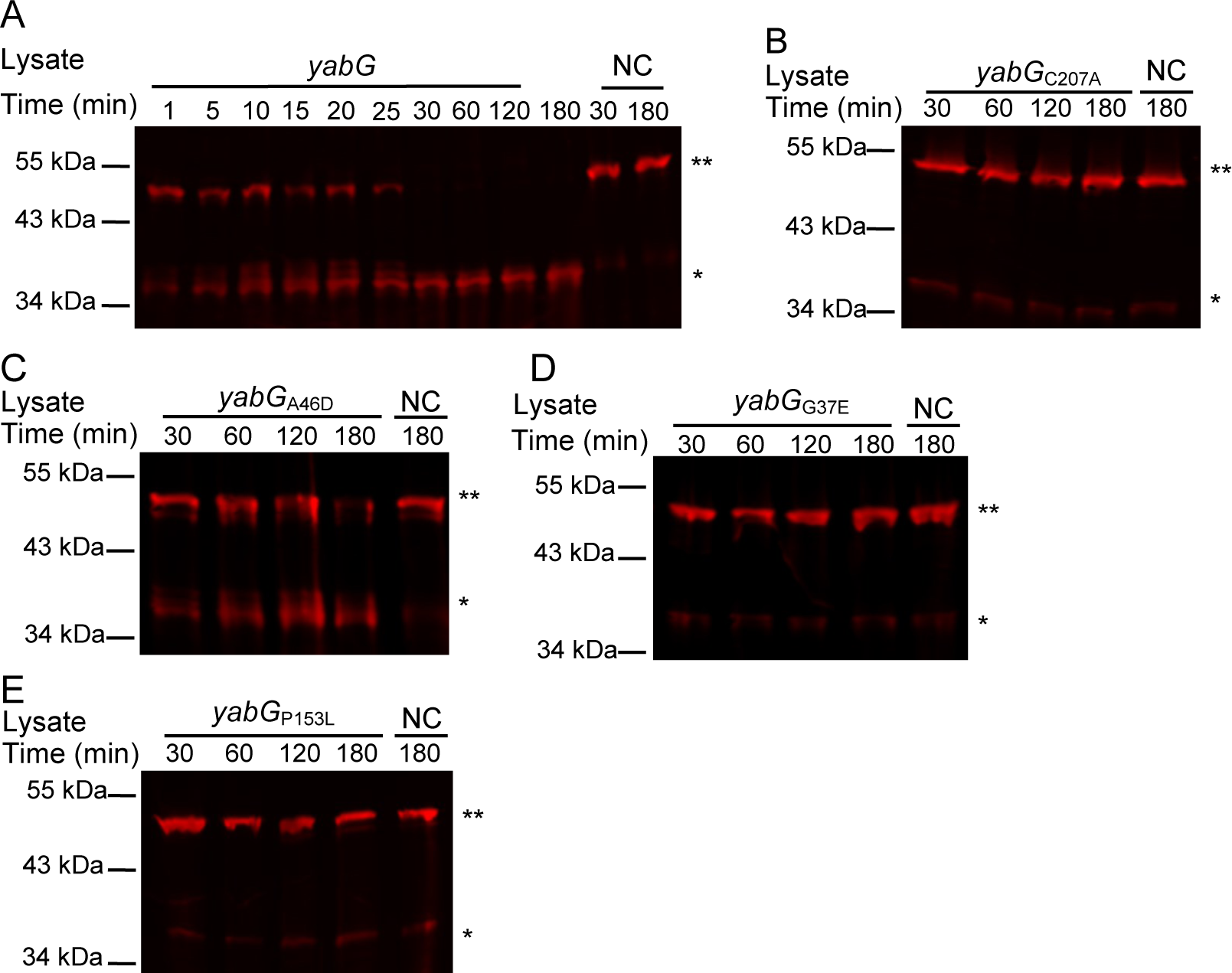
Processing of recombinant preproSleC by *yabG* alleles. Recombinantly expressed and purified preproSleC was incubated for the indicated times in *E. coli* lysate that expressed *yabG* allele (Table S3). A negative control sample of recombinant protein incubated in buffer was included (NC). Samples were boiled in SDS-sample buffer and separated by SDS-PAGE. SleC processing was analyzed using antisera specific for the SleC protein (the SleC antibody detects both preproSleC and proSleC forms). LI-COR Odyssey-CLx was then used to analyze the Western blots. A) Wild type *yabG* 1-180 minutes. B) *yabG*_C207A_ 30-180 minutes. C) *yabG*_A46D_ 30-180 minutes. D) *yabG*_P153L_ 30-180 minutes. E) *yabG*_G37E_ 30-180 minutes. ** indicates the pre-proSleC band and * indicates the proSleC band. Note: the purified SleC used in the assays with *yabG*_C207A_, *yabG*_G37E_, and *yabG*_P153L_ had proSleC present in the sample prior to the assay as seen in the controls.

**Supporting Figure 2.**
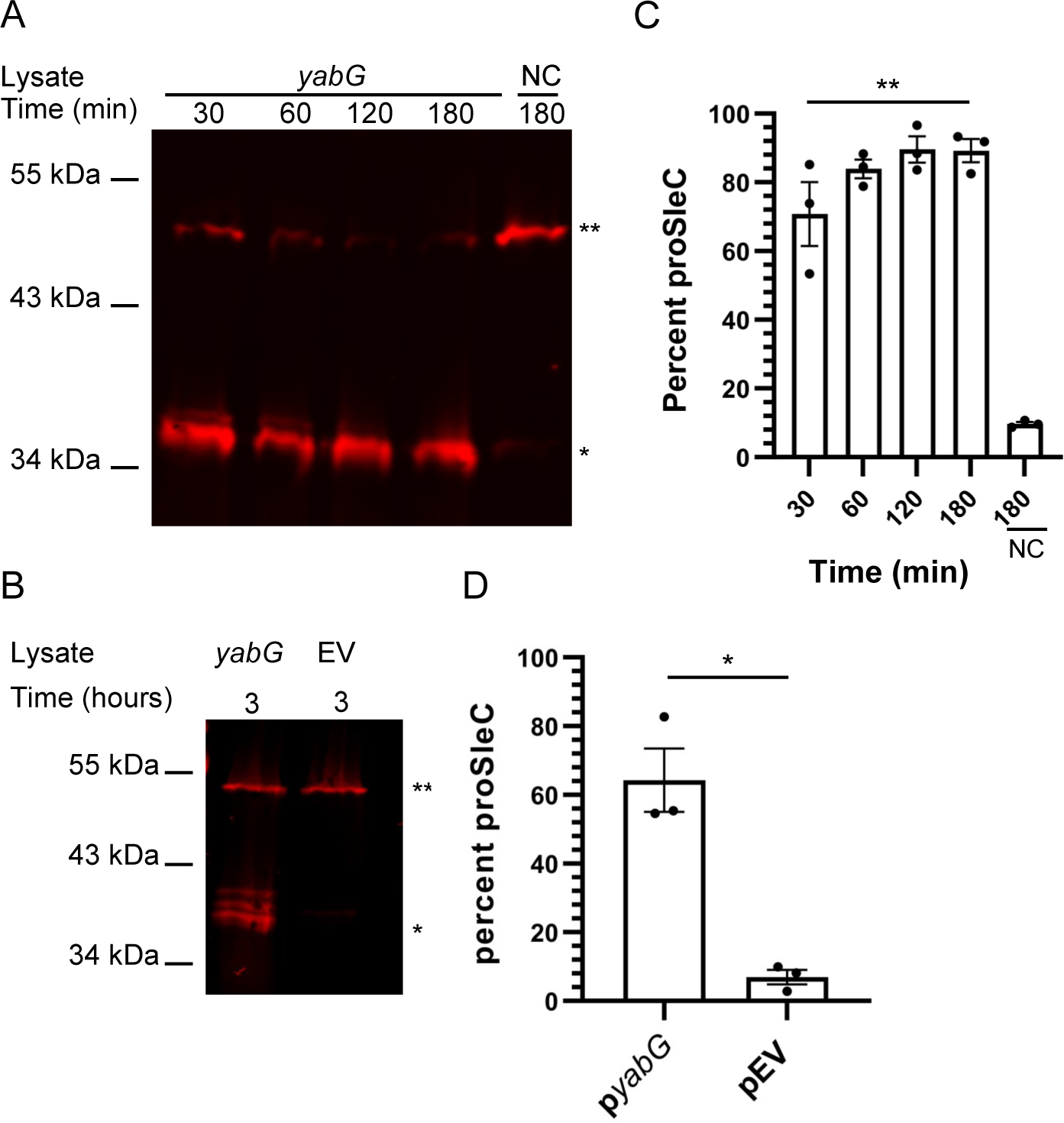
proSleC present in the purification samples does not affect YabG cleavage. Recombinantly expressed and purified preproSleC was incubated for the indicated times in *E. coli* lysate with / or without wildtype YabG. A control sample of recombinant protein incubated in buffer was included. Samples were boiled in SDS-sample buffer and separated by SDS-PAGE. SleC processing was analyzed using antisera specific for SleC protein (the SleC antibody detects both preproSleC and proSleC). LI-COR Odyssey-CLx was then used to analyze the Western blots. A) Wild type *yabG* 1-180 minutes Western blot. B) The percentage of processed proSleC from A). C) p*yabG* (pAC28) and pEV (pET22b) 180 minutes Western blots. A negative control sample of recombinant protein incubated in buffer was included (NC). D) Percentage of processed proSleC from B). The percentage of proSleC was quantified on the LI-COR Odyssey-CLx, [proSleC signal/(proSleC signal + preproSleC signal)]*100. ** indicates preproSleC and * indicates proSleC. The data represent the averages from three biological replicates and error bars represent the standard error of the mean. C) Statistical significance was determined using one-way ANOVA with Šídák’s multiple comparisons test (** < 0.0001). D) Statistical significance was determined using unpaired t-test (* <0.0021).

**Supporting Figure 3.**
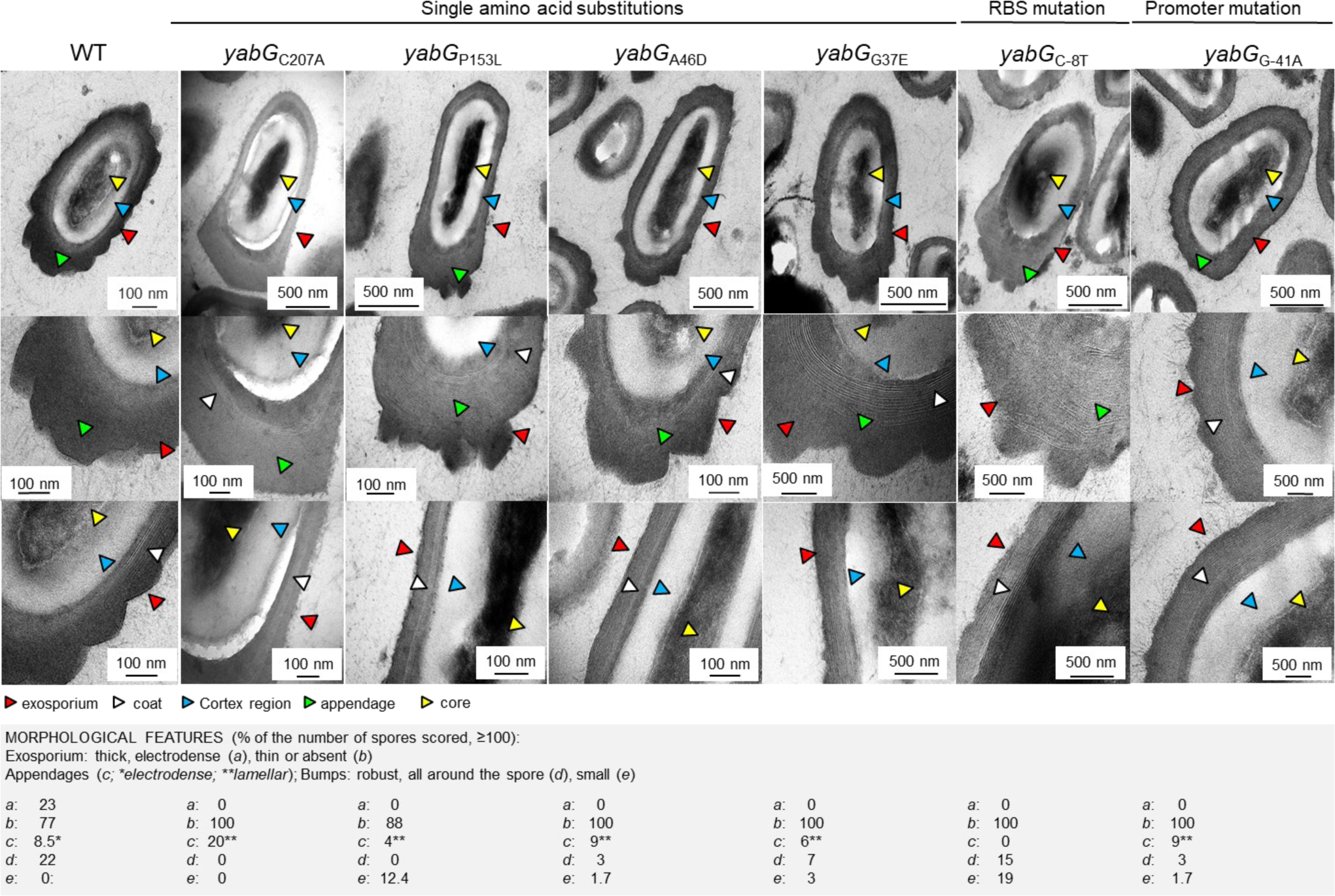
Morphological alteration in spores of the *yabG* mutants. Gradient purified spores of the WT (630Δ*erm*), *yabG*_C207A_ and the indicated point mutants were analyzed by thin sectioning transmission electron microscopy. The arrowheads point to: the edge of the exosporium, red; the coat, white; the cortex, blue; the spore core, yellow; the electrodense regions within the spore appendage, green. The table refer to the percentage of spores (> 100) in which the indicated feature is present; thick electrodense exosporium, a; thin or absent exosporium, b; electrodense lamellar appendage, c; bumps or robust appendage and all around the spore, d; and small bumpy appendage, e. Scale bars are indicated in the panels.

